# Accurate prediction of bacterial two-component signaling with a deep recurrent neural network *ORAKLE*

**DOI:** 10.1101/532721

**Authors:** Jan Balewski, Zachary F. Hallberg

## Abstract

Two-component systems (2CS) are a primary method that bacteria use to detect and respond to environmental stimuli. Receptor histidine kinases (HK) detect an environmental signal, activating the appropriate response regulator (RR). Genes for such *cognate* HK-RR pairs are often located proximally on the chromosome, allowing easier identification of the target for a particular signal. However, almost half of all HK and RR proteins are *orphans*, with no nearby partner, complicating identification of the proteins that respond to a particular signal. To address this problem, we trained a neural network on the amino acid sequences of known 2CS pairs. Next, we developed a recommender algorithm that ranks a set of HKs for an arbitrary fixed RR and arbitrary species whose amino acid sequences are known. The recommender strongly favors known 2CS pairs, and correctly selects orphan pairs in *Escherichia coli*. We expect that use of these results will permit rapid discovery of orphan HK-RR pairs.

## Introduction

Two-component systems (2CS) are a ubiquitous signaling paradigm in bacteria, and control the response to a variety of stimuli, including various nutrients, molecules used in cell-to-cell communication, and light [1,2]. These systems are composed of two protein partners, referred to as a *cognate* pair of proteins [1,2]. In an archetypal two-component system, a membrane-bound histidine kinase (HK) senses an extracellular signal through an N-terminal sensory domain. In response to the structural changes upon ligand binding, the C-terminal kinase domain is activated, leading to autophosphorylation at a conserved histidine residue. This autophosphorylation event then permits transfer of the phosphate group from the histidine residue of the kinase to a conserved aspartate residue on the N-terminus of a cognate response receiver regulator (RR) domain. The conformational change brought about by phosphorylation of the RR domain then leads to activation of a C-terminal output domain to modulate enzymatic or cellular function. Most RR domains possess auto-phosphatase activity, leading to a timed cessation of activation [3].

Many cognate pairs are located adjacent to one another in the genome, usually within the same operon [4,5]. This property of many cognate proteins has been previously used to determine the residues involved in cognate interactions [4,6]. More difficult is the identification of pairing partners for *orphan* HKs or RRs, which have no adjacent partner, to their cognate receptor or kinase, respectively. Traditionally, discovery of orphan HK or RR cognate pairs requires *phosphotransfer profiling* [7], which involves individual experimental tests between the orphan protein and each potential pairing partner. To resolve all possible HK-RR pairs for a species whose genome consists of *n* HK proteins and *m* RR proteins it requires up to *n* × *m* tests. This task can be daunting, since for some species there are up to 10^4^ possible 2CS pair combinations, such as in the deltaproteobacteria *Myxococcus xanthus* [8] and *Geobacter sulfurreducens* [9].

Early computational work to predict HK-RR cognates focused on statistical methods related to genome location [10]. A Bayesian method was developed to predict orphan HK-RR pairs from sequenced bacterial genomes [10]; in particular, this method does not require the residues important for the interaction to be known. However, the Bayesian method assumes that orphan kinases are always paired with orphan response regulators, and that there is a direct one-to-one correlation between kinase and response regulator, but exceptions to both of these assumptions exist [11,12]. Improved prediction algorithms use direct coupling analysis (DCA), in which multiple sequence alignments of known HK-RR pairs are used to generate scoring functions for a given kinase/response regulator pair [13]. The newest computational prediction algorithm, available online as MetaPred2CS [14], utilizes a support vector machine method to combine six different prediction methods to compute a score for any two given components in a 2CS.

However, despite these advances in computational prediction of two-component system interactions, to our knowledge only one orphan kinase has been successfully linked to a response regulator using these methods [11], and there remains a need for a simple method to determine potential cognate pairs. Deep Learning (DL) based recommendation systems offers a compelling alternative for orphan HK-RR discovery. DL models have the capacity to aggregate information from many known 2CS pairs for a large number of species and generalize this knowledge to offer predictions for new pairs of proteins based solely on their amino acid sequences. We used a Long Short Term Memory network (LSTM) [15] based Machine Learning (ML) model that scores the likelihood that two arbitrary HK-RR proteins interact. LSTM are type of a Recurrent Neural Network often used in ML to infer information from sequential data, e.g. for natural language processing. LSTM has also been used in biology for DNA-based analysis [16–18]. In this work the HK and RR amino acid sequences are fed into two double-layered LSTM branches. The LSTM outputs are then concatenated with the phylogenetic class category and further processed by four layers of a fully-connected neural network. The recommender algorithm developed from this model, termed ORphAn Kinase Lead Engine (*ORAKLE*) that ranks HKs based on their likelihood of pairing with a given RR.

## Results

*ORAKLE* was trained on the available body of known 2 · 10^4^ cognate and 5 · 10^5^ non-cognate 2CS pairs for over 1300 bacteria species [19]. More details about the input data curation, the model, and the training process are provided in ‘Materials and Methods’ section.

We will use the following notations. A neural network is a function

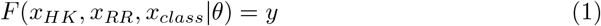

that accepts two sequences of amino acids: *x_HK_* (*x_RR_*) defines the HK (RR) proteins and *x_class_* refers to the phylogenetic class of the species. F also depends implicitly on the DL model parameters *θ*, which are determined during the training process. The output of the network, *ORAKLE score y*, is computed using the softmax function, which ensures that *y* satisfies *y* ∈ [0,1]. By convention, the higher the score of a pair, the more likely it forms a functional 2CS.

Any set of HK-RR pairs can be ordered by the *ORAKLE* score and indexed 0,1,…etc. This index is called *rank*. The pair with rank of 0 is the most likely to form 2CS, given the set of pairs.

For the purpose of this paper we have removed the 5 species listed in Table 1 from both training and validation sets. The *ORAKLE* predictions for the known 2CS pairs made for the species not used for training will be called *a priori*, while those for species used in the training will be called *a posteriori*.

**Table 1.**
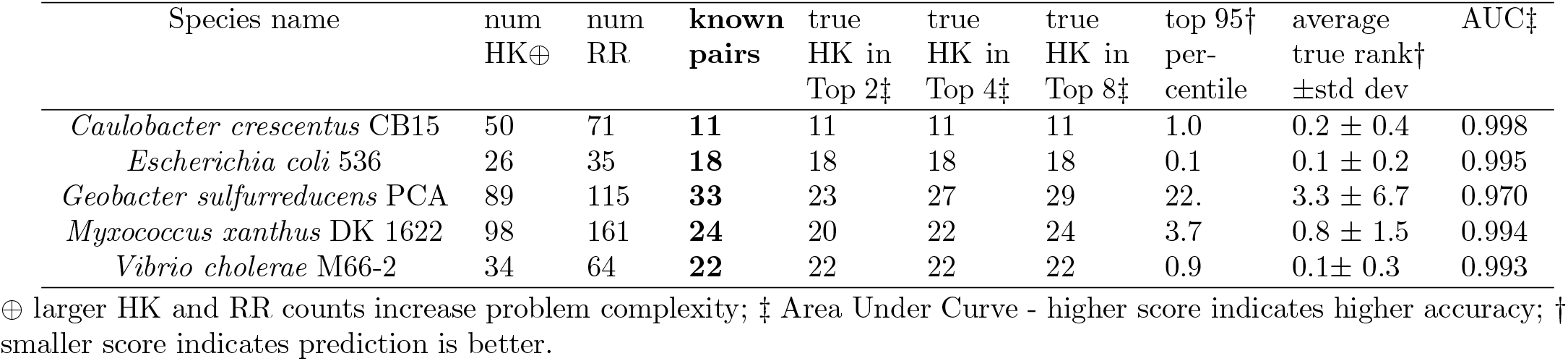
Accuracy metrics of *ORAKLE* prediction for selected species are shown in columns 5 to 10, see text for details. The 5 species listed in this table were excluded from any training and they are true *a priori* predictions.

We present several alternative metrics showcasing the ability for this neural network to pair histidine kinase-response regulator interactions. First, we demonstrate the accuracy of our *a posteriori* predictions for known kinase pairs for a subset of several hundred bacterial species out of 1300 used in the training. We then check the precision of *ORAKLE* by making *a priori* predictions for known 2CS pairs in *Escherichia Coli* and for 4 other species listed in Table 1.

### A *posteriori* Predictions for Gammaproteobacteria

We predicted the rank of the cognate HK for all RR proteins with a known interaction for 223 species from the Gammaproteobacteria class (Fig. 1). Under the assumption that orphan HK-RR pairs interact in a similar fashion, we expect this rank to be close to zero for orphan HKs/RRs, for which the interaction partner remains unknown.

**Fig 1.**
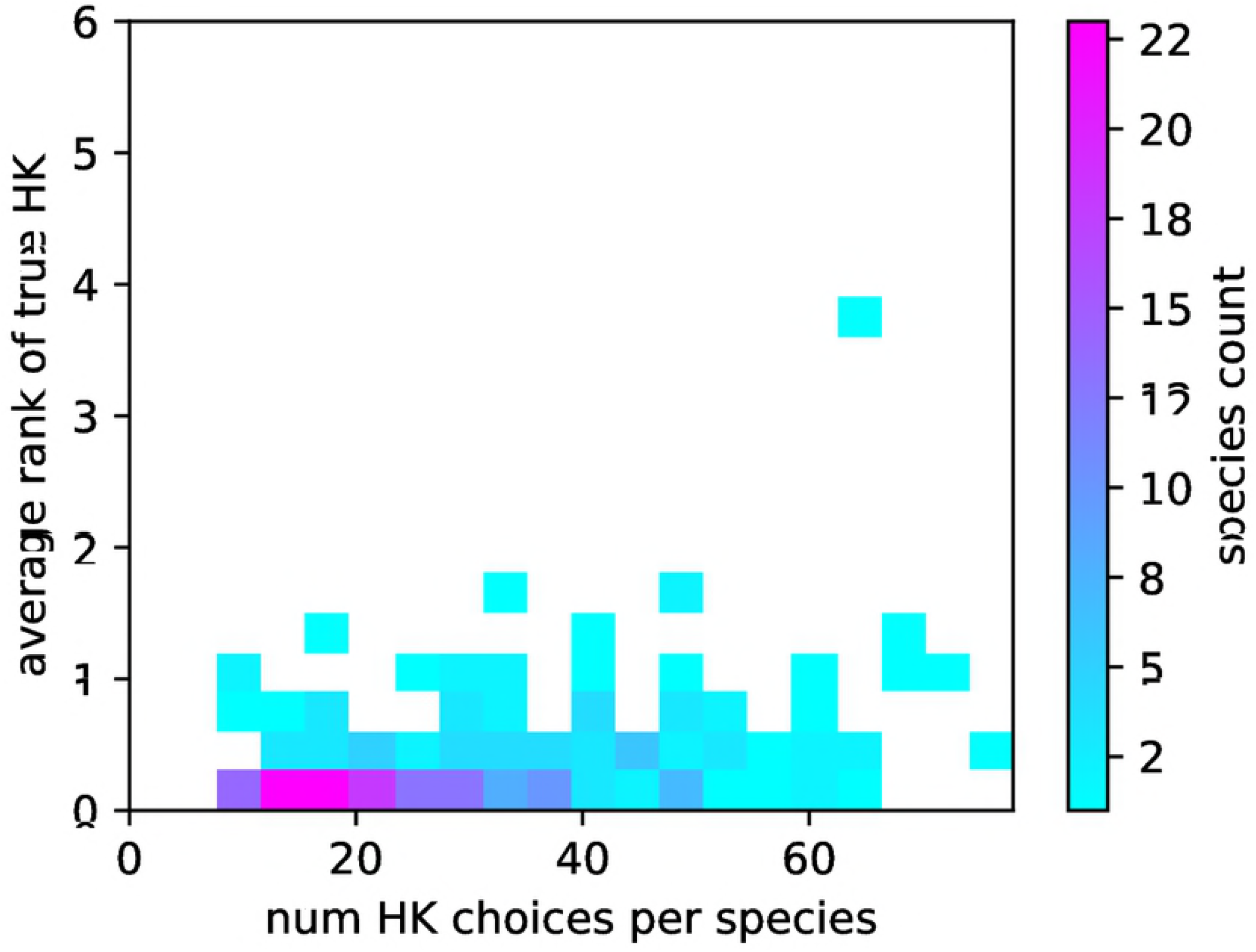
Distribution of the average rank (smaller is better) of the cognate HK per species for known cognate pairs for 223 species from the Gammaproteobacteria class. The number of HK (choices) per species is on the horizontal axis.

Despite the large number of HK choices in some species, and the presence of additional potential interactions that were not included in training (in the form of orphan kinases), we found that on average across 223 species, the correct HK was ranked as 0.3 ± 0.4 - often the top choice. In the worst case species, *Allochromatium vinosum* DSM 180 with 66 HKs, the average rank of the known cognate HK, given each RR, was 3.7 ± 7.1. The accurate ranking observed throughout the bacterial phylogeny (Table 2), demonstrating that 2CS can be readily paired using a neural-network based prediction algorithm. The worst prediction accuracy is observed for the phylogenetic ‘group’ labeled ‘others’^1^, whose class information was not used by the *ORAKLE*.

**Table 2.** Phylogenetic classes used for training. For each species, the average HK rank given RR was computed, and was averaged over the species in the class.

| phylogenetic class | species | known<br>pairs | average<br>true HK<br>rank ±std<br>dev |
| --- | --- | --- | --- |
| Actinobacteridae | 130 | 3305 | $0.3 \pm 0.3$ |
| Alphaproteobacteria | 129 | 1737 | $0.6 \pm 1.0$ |
| Bacillales | 94 | 1854 | $0.2 \pm 0.2$ |
| Betaproteobacteria | 102 | 2339 | $0.5 \pm 0.7$ |
| Clostridia | 111 | 2059 | $0.4 \pm 0.3$ |
| Cytophagia | 12 | 252 | $1.5 \pm 0.9$ |
| Deltaproteobacteria | 55 | 1277 | $1.4 \pm 1.2$ |
| Epsilonproteobacteria | 19 | 345 | $0.8 \pm 0.6$ |
| Flavobacteriia | 24 | 326 | $0.7 \pm 0.5$ |
| Gammaproteobacteria | 224 | 3965 | $0.3 \pm 0.4$ |
| Lactobacillales | 27 | 461 | $0.2 \pm 0.2$ |
| Sphingobacteriia | 8 | 221 | $1.2 \pm 0.7$ |
| Spirochaetales | 16 | 249 | $2.1 \pm 2.4$ |
| all others | 226 | 2378 | $2.2 \pm 3.0$ |

### A *priori* predictions for *E. coli*

With *a posteriori* predictions demonstrating that a neural network-based approach to predicting HK-RR interactions was feasible, we chose to investigate the *a priori* accuracy of predictions in a well-studied strain with known HK-RR pairs that was not included in the training set. The uropathogenic strain of *E. coli*, O6:K15:H31 536 [20], possesses a genome with 26 HK proteins and 35 RR proteins, of which 18 HK-RR pairs are confirmed based on proximity in the genome [19].

The phosphate regulation HK-RR pair PhoR/PhoB (corresponding to ECP_0459/ECP_0458) has been validated in a number of studies [21]. Indeed, we find that this pair is readily predicted by the *ORAKLE*, with both PhoR being given as the top choice to PhoB, and PhoB the top choice for PhoR (Fig. 2). Similarly, other known HK-RR pairs, such as the nitrate-sensing NarL/NarX (ECP_1270/ECP_1271) [22] and envelope stress-sensing BaeR/BaeS (ECP_2119/ECP_2118) [23], were readily matched *a priori* (Table 3). Only in one case for a known 2CS pair, ECP_1902/ECP_1901, which have unknown function, was the top choice incorrect.

**Fig 2.**
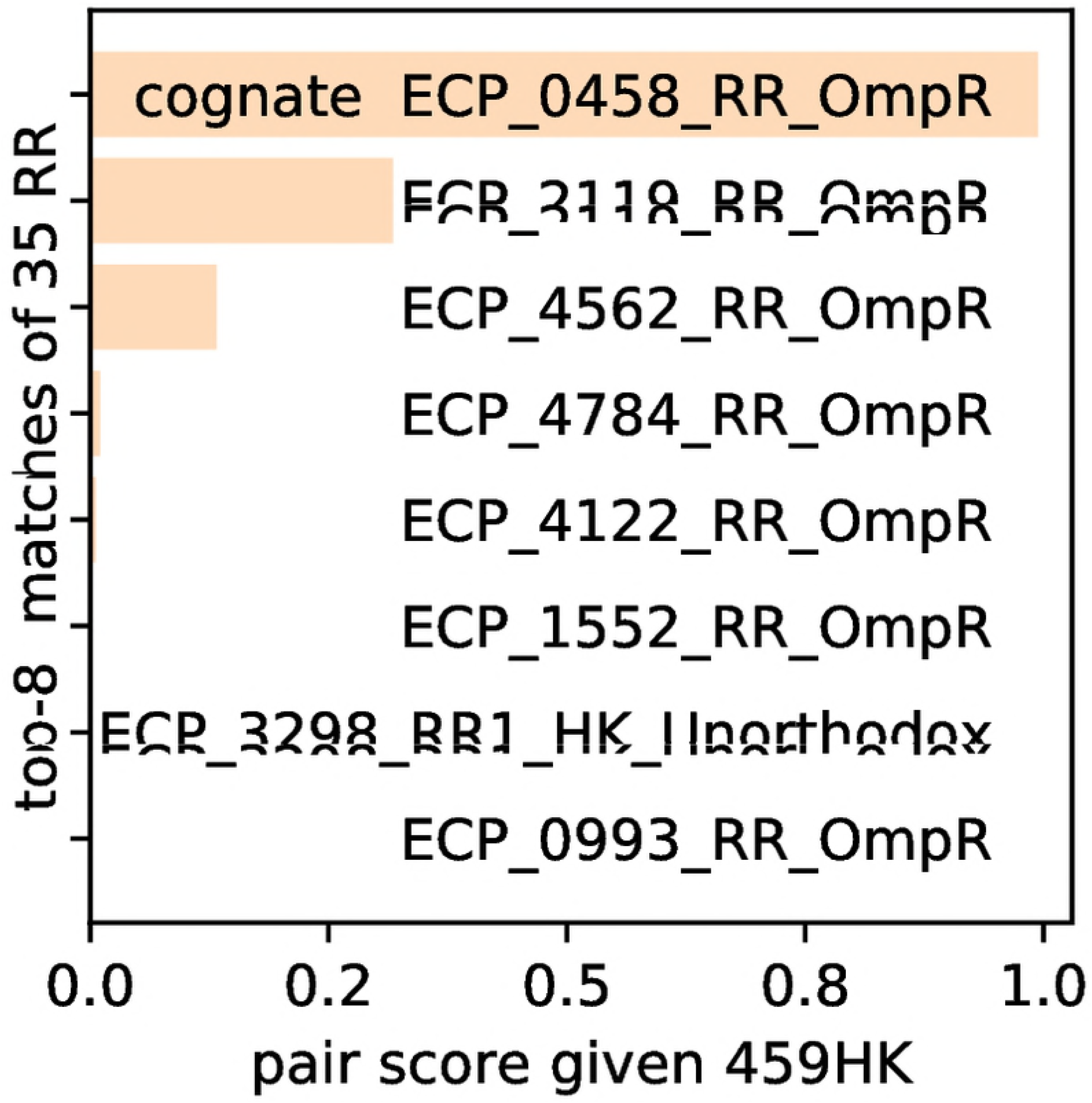
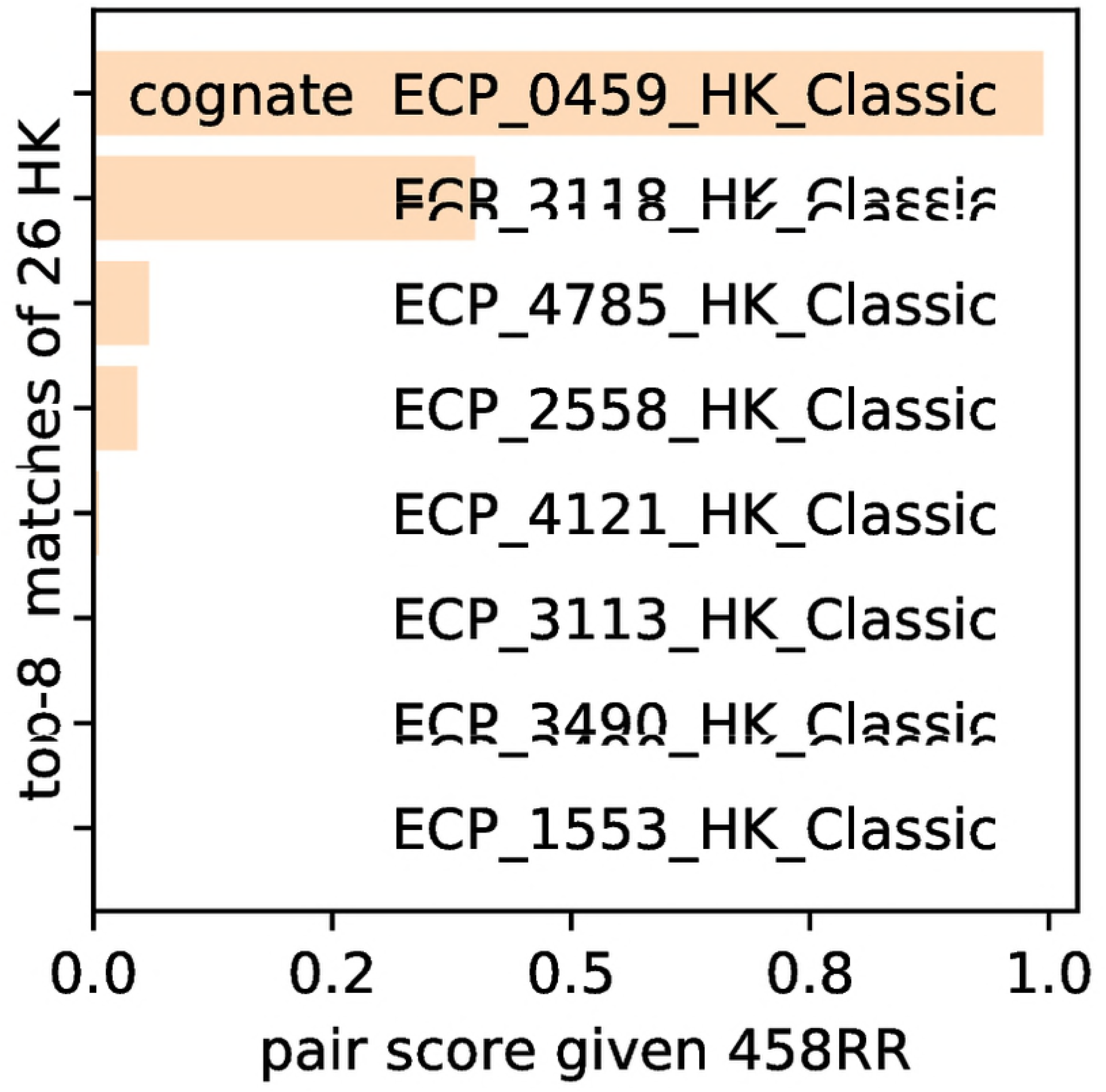
*A priori* predictions for *E. coli*. The known HK-RR pair PhoR/PhoB, corresponding to ECP0459/ECP0458, was correctly predicted as the highest ranked match for both approaches: Left) given 459HK scanned against all 35 RRs. Right) given 459RR scanned against all 26HKs. The magnitude of scores is denoted by the length of the horizontal bars.

**Table 3.** The rank of the cognate HK given a RR for unambiguous 2CS pairs in *E. coli*. 26 choices of HK were considered for every listed RR. Only the first 10 alphabetically sorted pairs are shown.

| given RR | true HK | rank ★ |
| --- | --- | --- |
| ECP_0458_RR_OmpR | ECP_0459_HK_Classic | 0 |
| ECP_0602_RR_OmpR | ECP_0601_HK_Classic | 0 |
| ECP_1125_RR_OmpR | ECP_1124_HK_Classic | 0 |
| ECP_1270_RR_NarL | ECP_1271_HK_Classic | 0 |
| ECP_1552_RR_OmpR | ECP_1553_HK_Classic | 0 |
| ECP_1902_RR_OmpR | ECP_1901_HK_Classic | 1 |
| ECP_2119_RR_OmpR | ECP_2118_HK_Classic | 0 |
| ECP_2163_RR_LytTR | ECP_2164_HK_Classic | 0 |
| ECP_2394_RR_NarL | ECP_2395_HK_Unorth | 0 |
| ECP_2975_RR_PrrA | ECP_2976_HK_Classic | 0 |
| ECP_3112_RR_OmpR | ECP_3113_HK_Classic | 0 |

★ predicted rank of true HK, rank 0 is the highest.

### Predictions for all known pairs of *E. coli*

There are many complementary metrics to determine the accuracy of *ORAKLE* predictions. We compare various metrics using the same set of 468 scores for the 18 RRs from the known cognate pairs against all 26 possible HKs of *E. coli* (Table 1). The highest-ranked HK (of 26 choices) is correctly identified for 17 out of 18 known RRs for *E. coli* (Table 3). The average rank of the cognate HK for known cognate HK-RR pairs in *E. coli* is 0.1 ± 0.3. The 95 percentile for the rank of the cognate HK is 0.1. If corresponding validation experiments were run on the top HK candidate for each of the 18 RRs, then one would likely fail. The distributions of scores for the 18 known cognate pairs and 450 non-cognate^2^ were calculated (Fig. 3). The initial signal to noise ratio (SNR) is 0.04. If we accept only pairs with a score above 0.9, we retain all 18 cognate pairs, i.e. the true positive rate is 1.0. The false positive rate is 0.02 in this situation. With this scoring metric, we keep 9 non-cognate pairs. Hence *ORAKLE* would improve SNR for *E. coli* 50 times to 2.0. The *Receiver operating characteristic* curve for *E. coli* (Fig. 4) approaches a perfect discriminator, since its area under the curve (AUC) is of 0.99.

**Fig 3.**
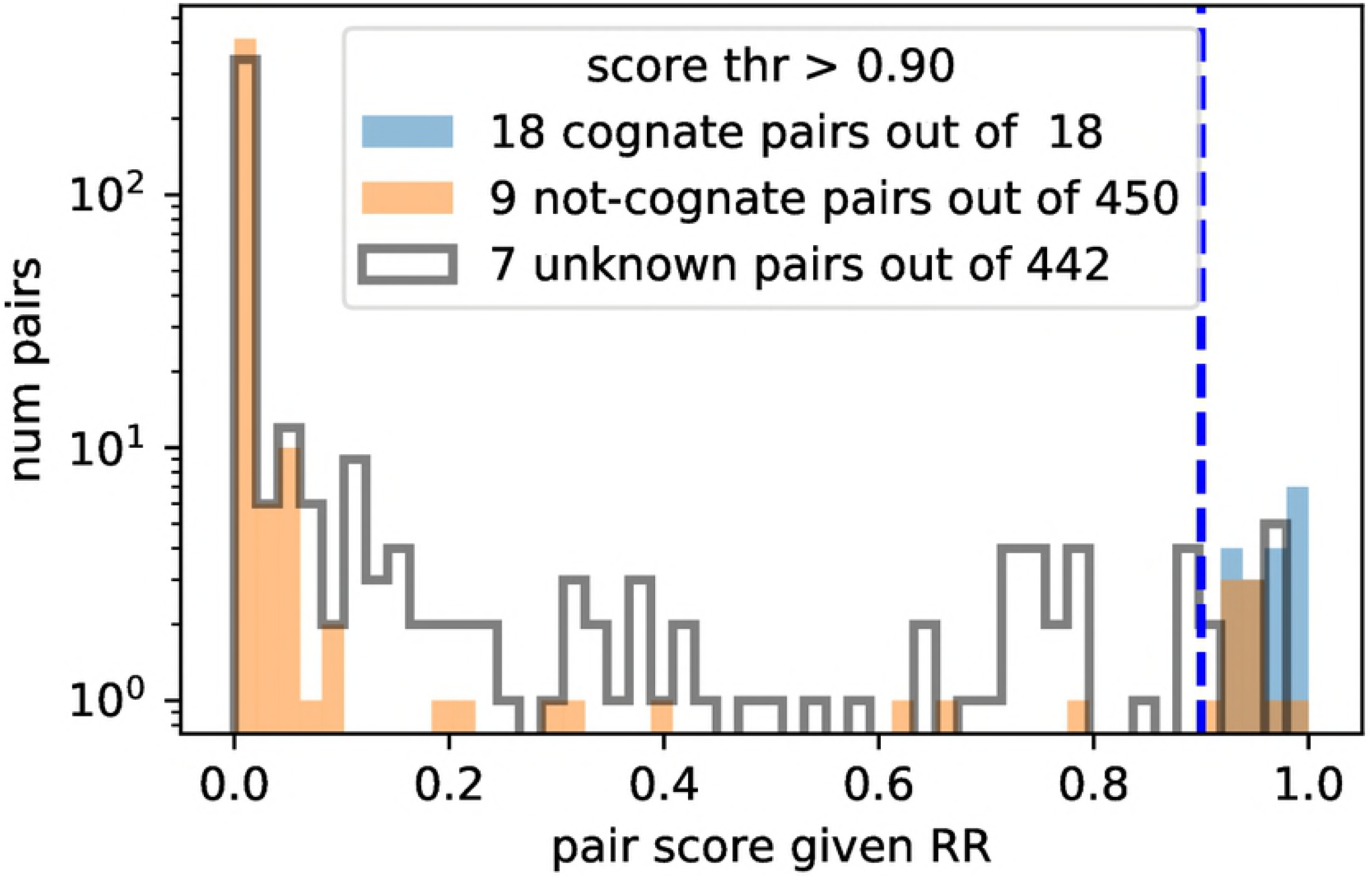
Distributions of scores for *E. coli* for known cognate pairs (orange), known non-cognate pairs (blue), and unknown pairs (black). The numbers in the legend show the count of pairs with the score above 0.9 (vertical dashed line).

**Fig 4.**
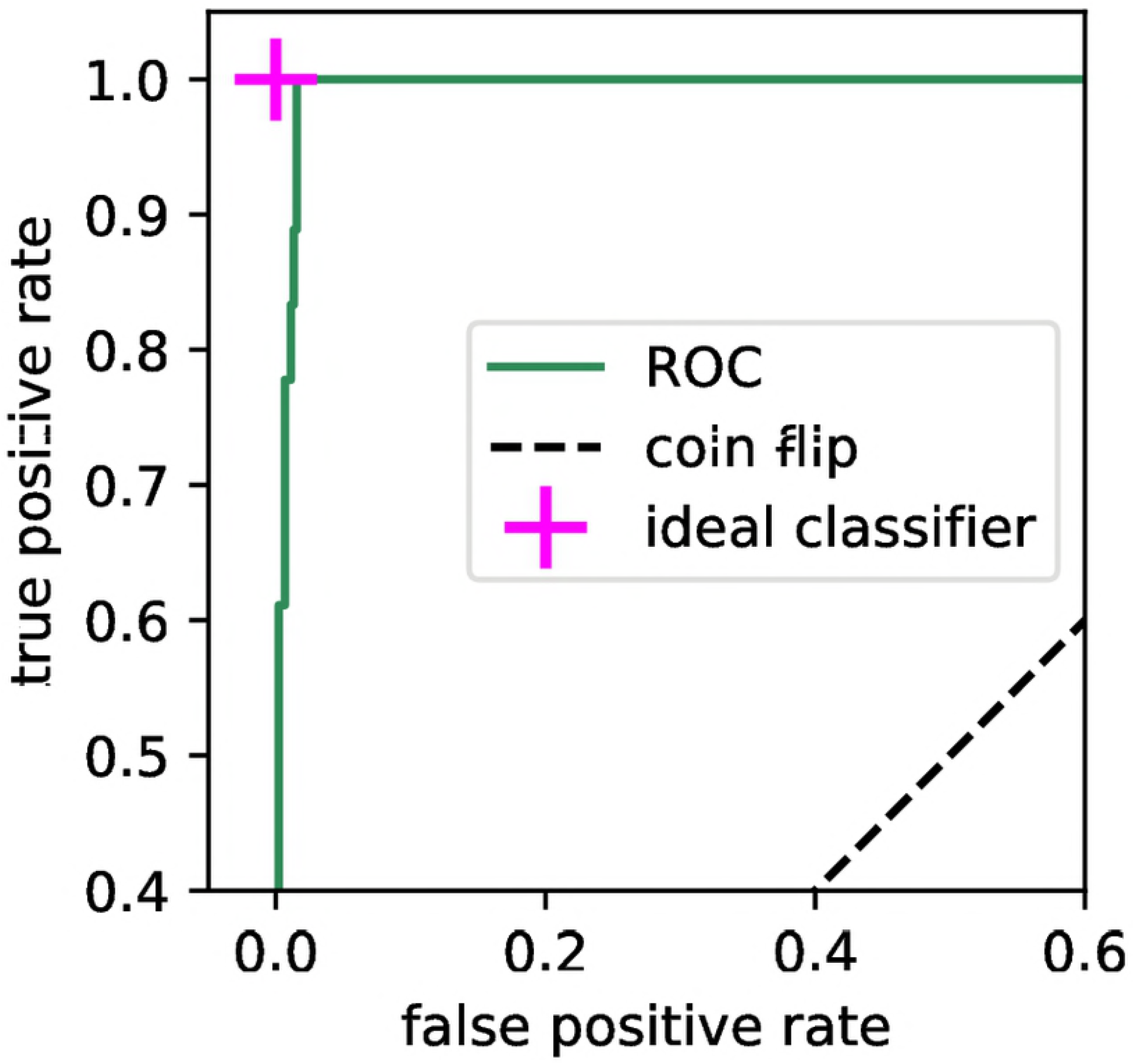
Receiver Operating Characteristic (ROC) curve for *E. coli* has an area of 0.995. Here cognate pairs were not oversampled.

To demonstrate the full potential of the *ORAKLE* we computed the scores for all 910 possible HK and RR pairs of *E. coli*, selected the top-2 HKs based on the score (Fig. 5). No correct pairs are missed with our prediction algorithm.

**Fig 5.**
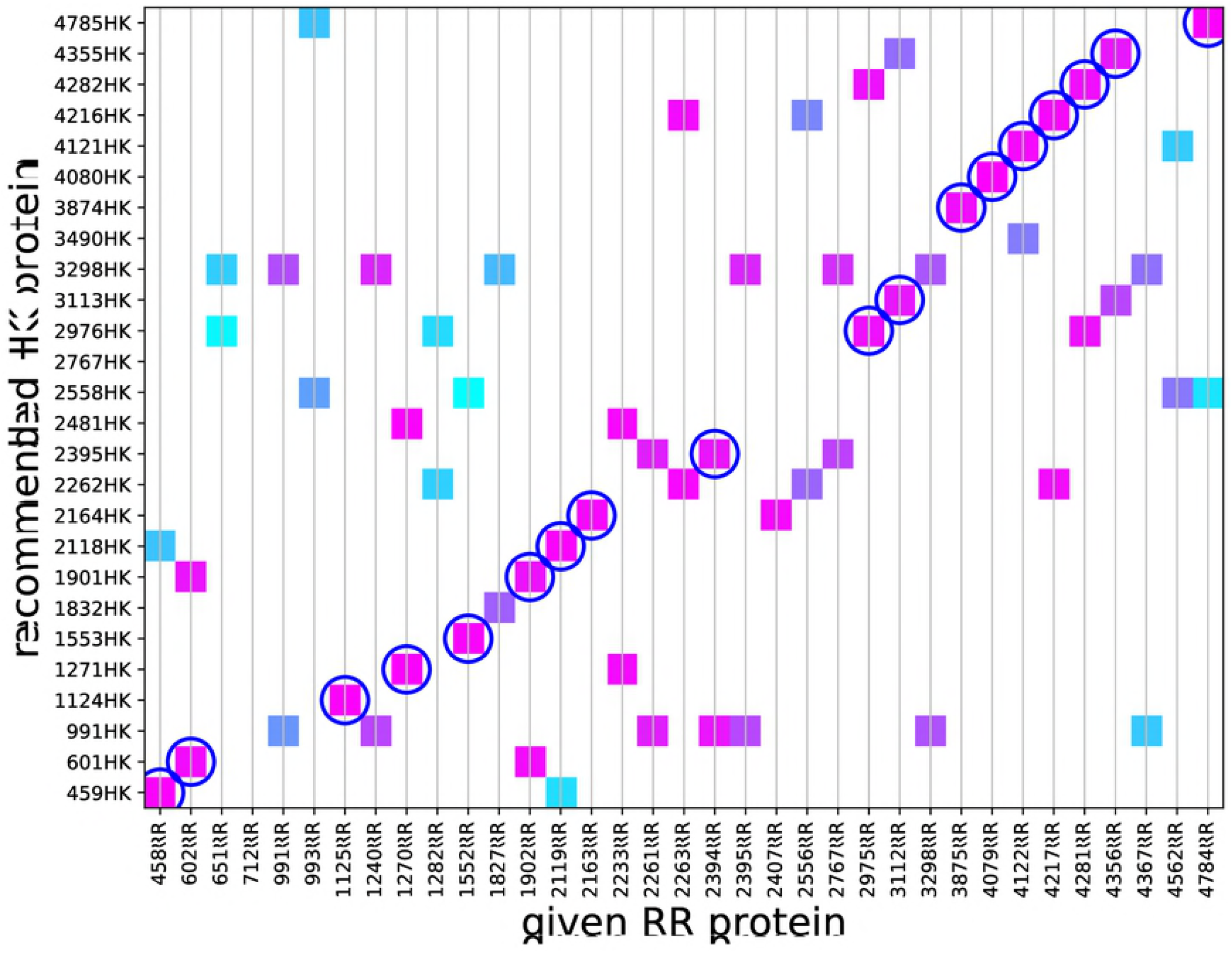
Score heat-map for the top-2 selection of HKs for each of 35 RRs for *E. coli*. The RRs listed on horizontal axis are ordered according to the location on the genome. The HKs are listed on the vertical axis, also order by location. Blue circles mark 18 known 2CS pairs of which all have been found.

### Recommendations for orphan proteins of *E. coli*

There are 17 orphan RR proteins in *E. coli* for which the sensor HK cannot be determined by proximity alone. We predicted the top 8 HK candidates for the orphan CheY response regulator in *E. coli*, involved in chemotaxis (Fig. 6). This orphan RR is a known target of the master chemotaxis regulator CheA (ECP_1832) [24]. Indeed, *ORAKLE* correctly predicted that CheA is the congate kinase for CheY, giving this kinase the only score above 0.5.

**Fig 6.**
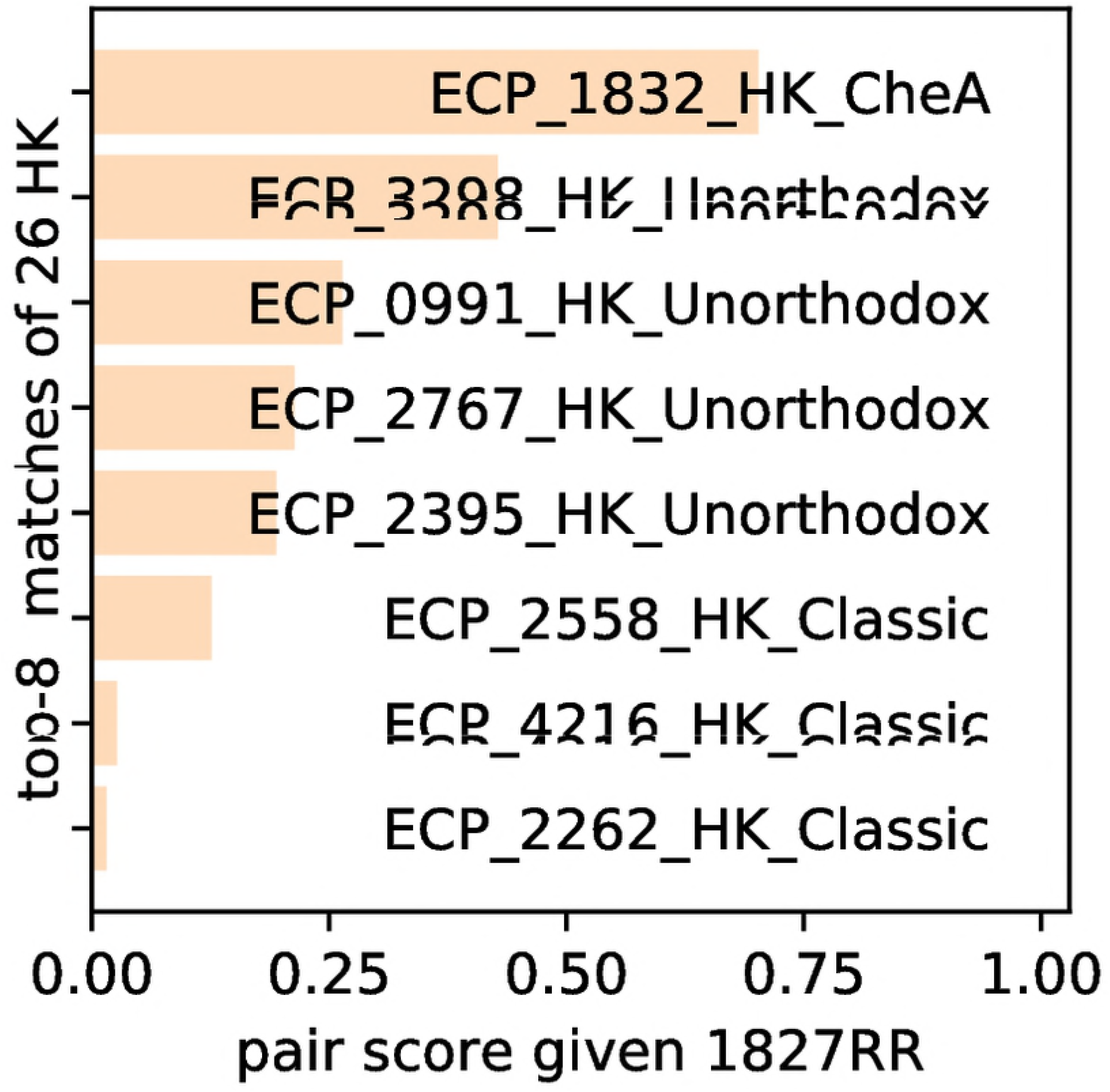
The top-8 most probable HKs for the *Orphan* 1827RR of *E. coli*

We used *ORAKLE* to score all 442 ‘unknown’ HK-RR pairs, including the 17 orphan RRs and the 26 *E. coli* HKs. The distribution of the score is shown in Fig. 3 as grey histogram. Only seven of those unknown pairs have a score above 0.9 (Table 4). Some of these predicted pairs are likely false positives. For instance, ECP_2261 is a hybrid histidine kinase-response regulator protein, consisting of an HK domain linked to a RR. RR domains in hybrid HK-RRs act as phosphate sinks, and do not obey the normal rules biochemical rules for pairing because they are tethered [25]. However, the predicted interactions between ECP_2395 and ECP_0991, may still point to positive interactions.

**Table 4.** Recommended HK matches for orphan RRs for *E. coli*, selected based on *ORAKLE* score above 0.9. The full distribution of scores is shown in Fig. 3.

| given RR | predicted HK | score |
| --- | --- | --- |
| ECP_2233_RR_NarL | ECP_2481_HK_Classic | 0.973 |
| ECP_2233_RR_NarL | ECP_1271_HK_Classic | 0.966 |
| ECP_2261_RR1_HK | ECP_2395_HK_Unorth | 0.904 |
| ECP_2261_RR1_HK | ECP_0991_HK_Unorth | 0.904 |
| ECP_2263_RR_NtrC | ECP_2262_HK_Classic | 0.974 |
| ECP_2263_RR_NtrC | ECP_4216_HK_Classic | 0.964 |
| ECP_2407_RR_Lyt | ECP_2164_HK_Classic | 0.968 |

The *ORAKLE* prediction can also differentiate between two potential response regulators for which assignment of the cognate kinase is ambiguous (Table 4). The kinase ECP_2262 is located in close proximity to both ECP_2261 and ECP_2263, preventing rapid assignment of its correct pairing partner. Indeed, *ORAKLE* assigns ECP_2263 as the correct partner.

### A priori predictions for other species

The same accuracy metrics were computed for the 4 other strains exluded from training (Table 1). Predictions of the *ORAKLE* remain fairly accurate even for species that contain a larger number of proteins involved in 2CS, and thus a much higher potential pair count e.g. *Myxococcus xanthus* or *Geobacter sulfurreducens*. In both these cases, known cognate pairs remain identified with a high accuracy even for more complex species.

With this prediction algorithm, we can generate a rank list for any set of possible RR-HK pairs for any bacterial species, given the amino acid sequences of these proteins (Table 1).

## Discussion

2CS are pervasive in the bacterial phylogeny, with most species having at least one and often dozens of genes encoding HKs and RRs. However, while these systems do not feature the crosstalk so widely seen in eukaryotic signaling systems [33], identification of isolated signaling networks remains difficult because orphan HKs and RRs, approximately half of all HKs and RRs, scattered throughout genomes do not have readily identifiable partners.

HK-RR pairs can be identified through many methods with varying accuracy, such as pooled transposon screening [34] and phosphotransfer profiling [7]. However, the former two methods require a condition under which the known member of the pair is essential for growth, and many HKs and RRs have no known functional role. Additionally, phosphotransfer profiling is often laborious, especially for large genomes. We expect that the *ORAKLE* will facilitate the rapid assignment of orphan HK-RR interactions, and provide a complementary ranking compared to previous orphan predictors [10,14].

Assuming that orphan HK-RR interactions are similar to those of neighboring pairs, the pairs sorted by the score from *ORAKLE* will favor the correct pair on the 2nd ±1 position on average [40], so users need only choose the top 4 HKs for a given RR (or top 4 RRs for a given HK) to have a high chance chance of testing the correct pairing member — an order-of-magnitude improvement over the use of brute-force methodologies. In practice, we find that the pairing partner is often scored by *ORAKLE* above 0.5. Thus, it appears that RRs that have no HK with a score above 0.5 are candidate noncanonical RRs, which may be activated through alternative mechanisms such as small molecule binding [36], serine/threonine kinases [35], or more complex phosphorelays [37].

While predictions of our algorithm are fairly accurate for known pairs, addition of more in vitro experimental results into the training set could further improve the predictive power of the *ORAKLE*. Currently we assume binary classification of HK-RR interactions during the training phase; the correct matches are given a score of 1, while unpaired/incorrect interactions are given a score of 0. This assumption ignores the wide multitude of crosstalk often observed between these proteins in vitro. A better training metric would be a continuous, experimentally-determined strength of interactions between all HKs and RRs from one species (such as Myxococcus xanthus). Further improvement could include also the spatial location of (or distance between) proteins, as some HK-RRs are spatially segregated into signaling compartments [38], which can act to reduce potential crosstalk. However, at this time we do not have sufficient experimental data to include in the training.

In the future, the *ORAKLE* training could also include the identity of the species which would extend its utility also for synthetic biology applications. In many cases, importing HK/RR pairs from other genomes have been used to create environmental sensors [39]. However, this application requires judicious selection of the host bacterium and of the HK/RR pair to be used, as crosstalk between the endogenous 2CSs present in the organism may hamper proper signaling. Such whole-species trained *ORAKLE* would make predictions which include impact from the crosstalk in the host bacterium and hence aid generation of new engineered kinases.

## Materials and Methods

### Input dataset

The P2CS database [19] is a comprehensive resource for the analysis of Prokaryotic Two-Component Systems. We have downloaded from there the sequences of all known HK and RR proteins in the fasta format, using P2CS version 4.5 from 2015. P2CS labels every protein as one of the following: ‘Orphan’, ‘Pair’, ‘Triad’, ‘Tetrad’, etc. We have used only those HKs or RRs labeled as ‘Pair’ for training, because only these HKs and RRs are uniquely bound. For all other proteins, only post-training predictions are made. We have removed duplicated sequences from the original P2CS database ^3^ and imposed self-consistency on HK-RR pairs, which lead to a training set of 1606 species with 2 · 10^4^ unique cognate HK-RR pairs. The average count of HK (RR) proteins per species was of 22 (34), the average cognate pair count per species was of 13, see Fig. 7 for the respective distributions.

**Fig 7.**
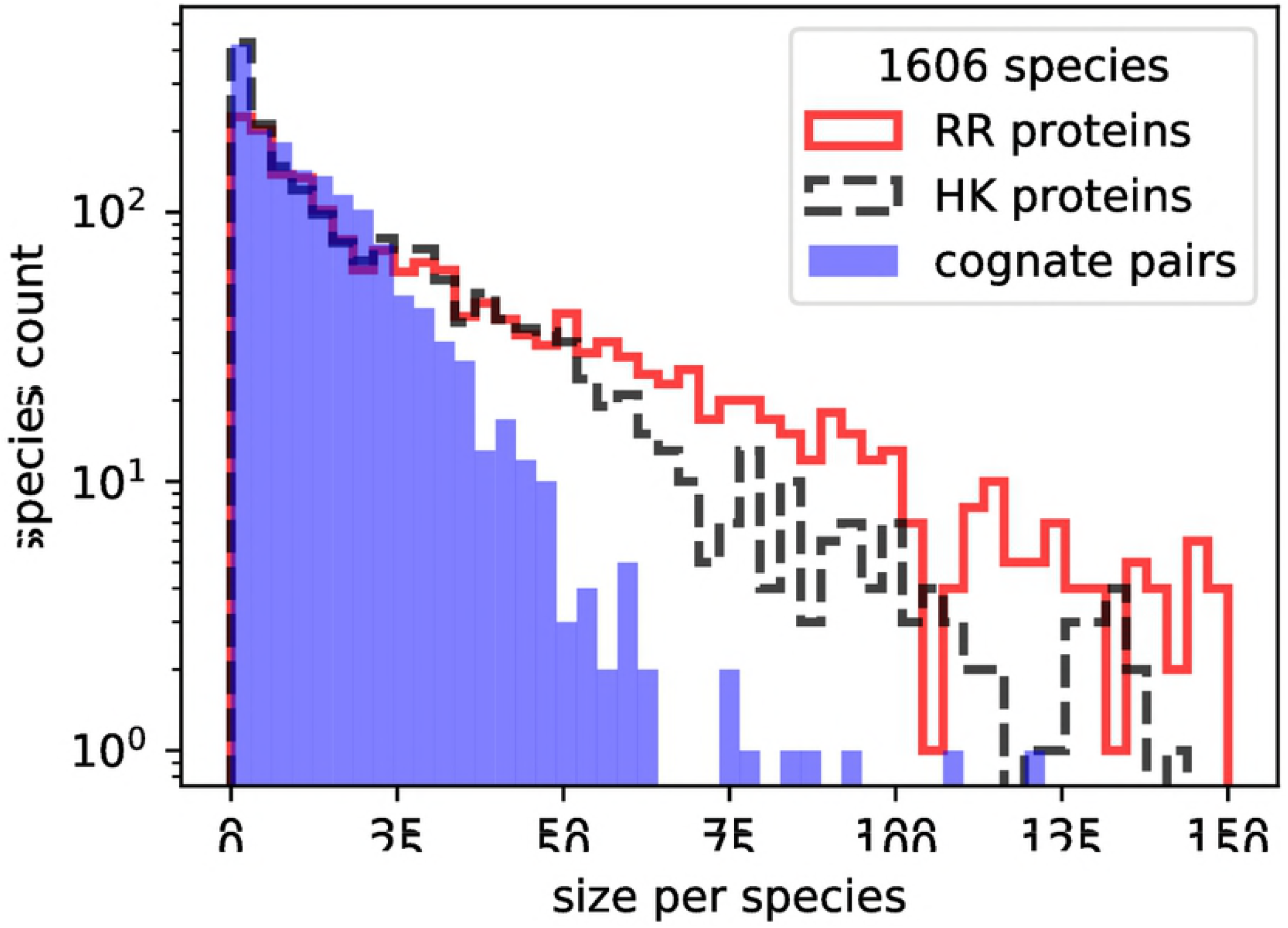
Number of HK, RR, and cognate pairs per species used for the training of the model. The data were obtained from P2CS database [19].

To inform the model about the differences between species we decided to use the phylogenetic class as an additional categorical feature. We have singled out the 13 most populous classes, listed in Table 2, out of 81 from the bacterial phylogeny. All remaining classes were merged into the ‘other’ class.

#### Negative examples

The supervised training method requires both positive and negative examples. We constructed the not-cognate pairs by permuting within each species the HK and RR proteins from the *N_cogn_* cognate pairs. Therefore, there are *N_cogn_*(*N_cogn_* − 1) known not-cognate HK-RR pairs per species - those are the negative training examples. To balance the ratio of the negative/positive pairs within each species we oversampled the minority cognate pairs by the factor of (*N_cogn_* − 1) and shuffled all pairs.

#### Formating protein sequences

The raw input amino acid sequences are of variable lengths, as shown in Fig. 8. We set-up the LSTMs [15] to expected a fixed length inputs. Therefore, we clipped too long HK (RR) proteins at 90 (130) amino acids or pre-padded with the 0-symbol the too short once, as needed.

**Fig 8.**
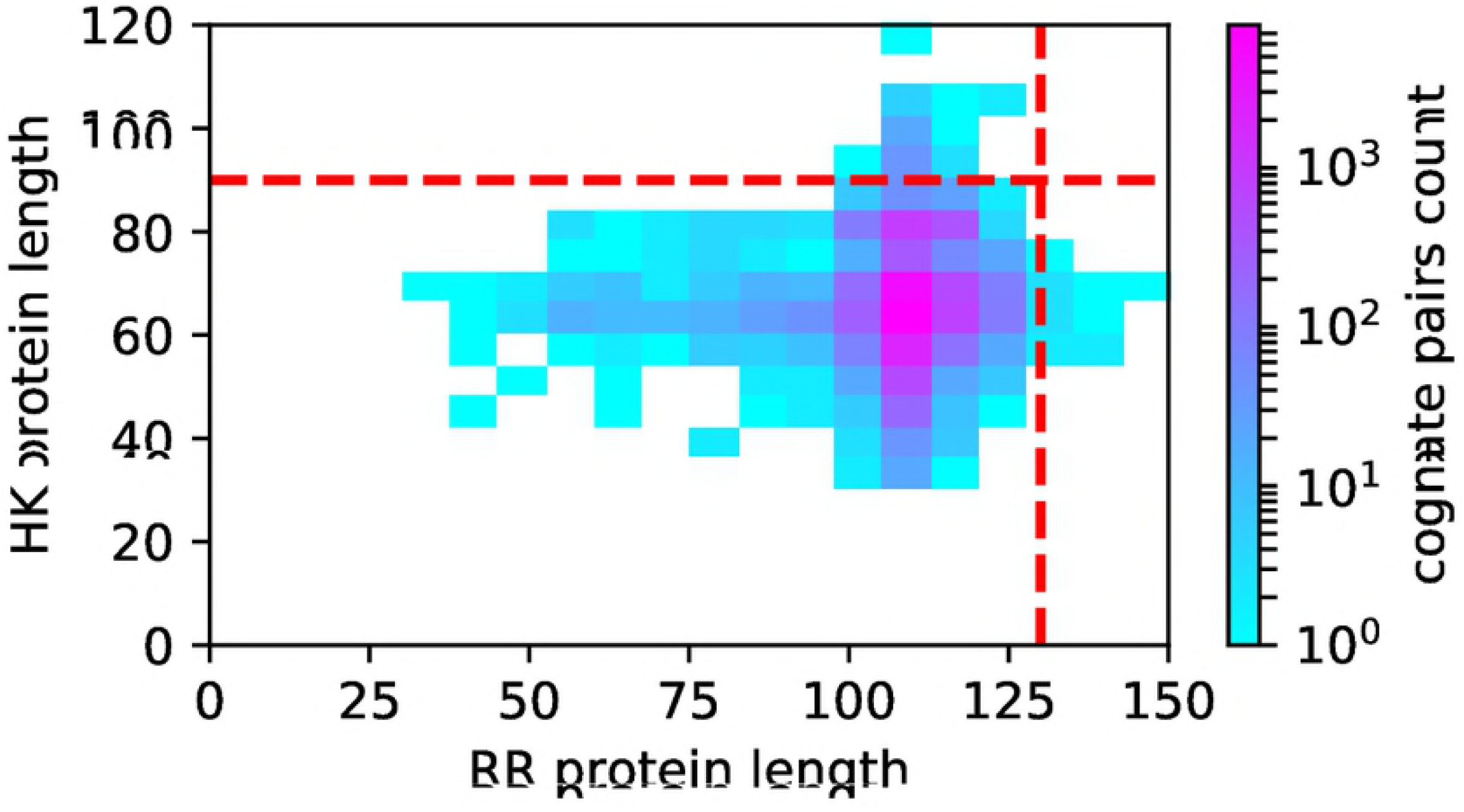
Correlation between the lengths of RR vs. HK amino acids for the known cognate pairs used in the training. The red dashed lines mark the clip value for respective type of proteins, see ‘Materials and Methods’ section for the details.

We have diversified the highly replicated cognate pairs at the initialization phase by randomly masking out 10% of amino acids in every input sequence and replacing their codes with the same 0-symbol used for the padding. This 10% noise was applied uniformly also to input sequences for the not-cognate pairs.

The 10% noise is an extension of the regularization, acting in a similar fashion as a dropout. Additionally, it insulates the model from possible systematic differences in accuracy in the protein sequences themselves, acquired due to different experimental methods used to assemble them. Most of modern gene sequencing/assembling methods make mistakes below 1% [26,27], which is negligible if added in quadrature with our induced noise of 10%.

All 3 inputs to the training: HK sequence, RR sequence, and phylogenetic class are categorical of size 22, 22, and 13, respectively. We 1-hot encoded all 3 categorical features. The final dimensions of the ML inputs are of 90 × 22, 130 × 22, and 1 × 13, respectively. Training data were shuffled before training commenced.

### Model topology

Since input data are sequences, use of the LSTM [15] unit was the most natural approach to convert the fixed length sequence into a one-dimensional feature vector. Our model, shown in Fig. 9, features two branches of double LSTM layers accepting amino acid sequences for HK and RR proteins. The resulting 65 and 110 features vectors are concatenated with the phylogenetic category input and passed to 4 fully connected layers that produce one score (Eq. 1). The dropout of 0.2 was applied between any layers, the LSTM recurrent dropout was set at 0.1, the random input noise was of 10%, as discussed earlier.

**Fig 9.**
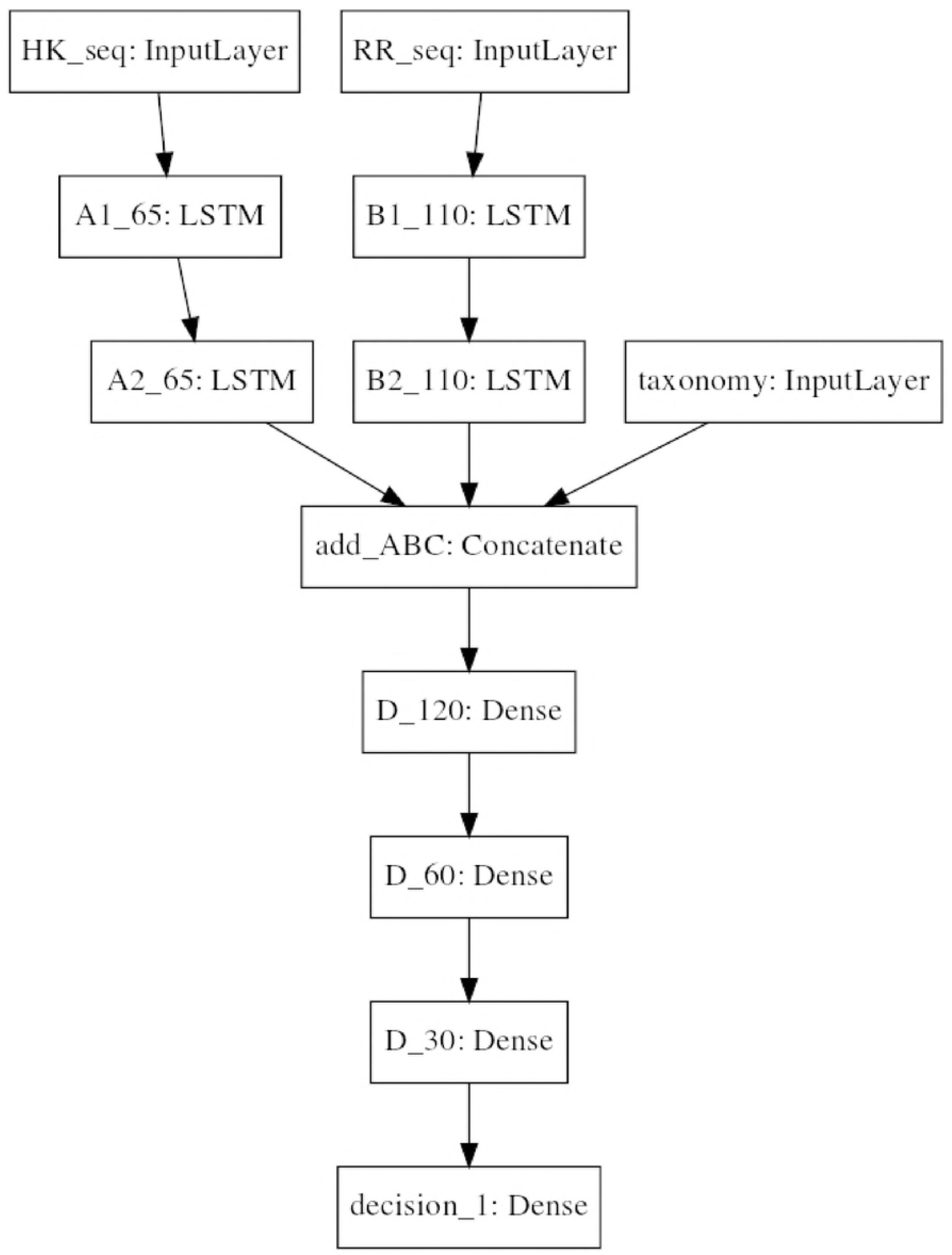
Model for *ORAKLE* Neural Network. Our model consists of two double LSTM layers whose outputs are concatenated with the phylogenetic class feature and further reduced to the single softmax score by 4 fully connected layers. The numbers in the boxes refer to the sizes of the output feature vectors. This model has the total of 243,801 parameters. The Dropout layers, not shown, are placed in-between all described layers.

This model was implemented in Tensorflow 1.3.0 [28] and Keras [29].

### Tuning of hyper-parameters

The final model, shown in Fig. 9, resulted from the evolution of the initial guess, followed by exploration of the hyper-parameter space as follows:

- The dropout rate between layers was varied between 0.05 to 0.5, leading to a optimal dropout of 0.2.
- The recurrent dropout rate inside LSTM was varied between 0.05 to 0.15, with a final recurrent dropout rate of 0.1.
- The dropout on input was varied between 0.05-0.40 to mitigate loss variability caused by the oversampling, with a final dropout on input of 0.1.
- The model size was varied between 14k and 550k parameters by proportional scaling of the number of features in all layers yielded up to 10% changes in AUC, with a final size of 240k parameters.
- Addition of a reduced learning rate on plateau by a factor of 0.3 every 3 epochs lead to improved AUC.
- Addition of more dense or LSTM layers was not beneficial.
- A change from oversampling of the minority class to sample weighting and preserving the natural mixture of labels of 1/(*N_cogn_* − 1) lead to more erratic convergence and was abandoned.
- A change from over-sampling of the minority class to under-sampling of the majority class lead to lower AUC and was abandoned.
- The minimizer finds different local minima even if the same input data are used due to a different starting seed, see Fig. 13 discussed later. We decided to train twice on each of 10 k-folds and average the scores over 20 models *θ_k_* to obtain higher accuracy of the averaged predicted score.

It is worth noting that there was no obvious optimal configuration of the hyper-parameters. Rather, the accuracy of prediction or stability of the training degrades if the working point moves away from the chosen one. Finally, the cost of exploring the hyper-parameter space is too large to sample all possible initial conditions. Our base-line model has a very high accuracy, with an AUC of 97%. For each configuration, a full training of approx. 1 CPU-year was required to reach a sub-percent statistical accuracy of AUC, to verify a new model’s accuracy.

### Model training

The model was trained with a *binary cross-entropy* loss function [30] and *Adam* optimizer [31]. We performed supervised learning on the balanced set of 5 · 10^5^ pairs with a batch size of 128. The initial learning rate was set to 0.001.

We used the K-fold cross-validation method, setting K=10, with 8 data segments merged as the ‘training’ set, 1 validation segment that provided only the loss as a feedback during the training, and 1 ‘test’ segment that was ignored during the training. We performed 10 independent trainings, cycling the segments to allow each of 10 segments to influence 8 models *θ_k_*. It required 19 GB of RAM to keep all 1-hot encoded input data in the memory. A single training required about an hour per epoch on an Intel Xeon Processor E5-2698, ‘Haswell’, node with 32 physical cores. The model convergence was reached after about 20 epochs, as shown in Fig. 10.

**Fig 10.**
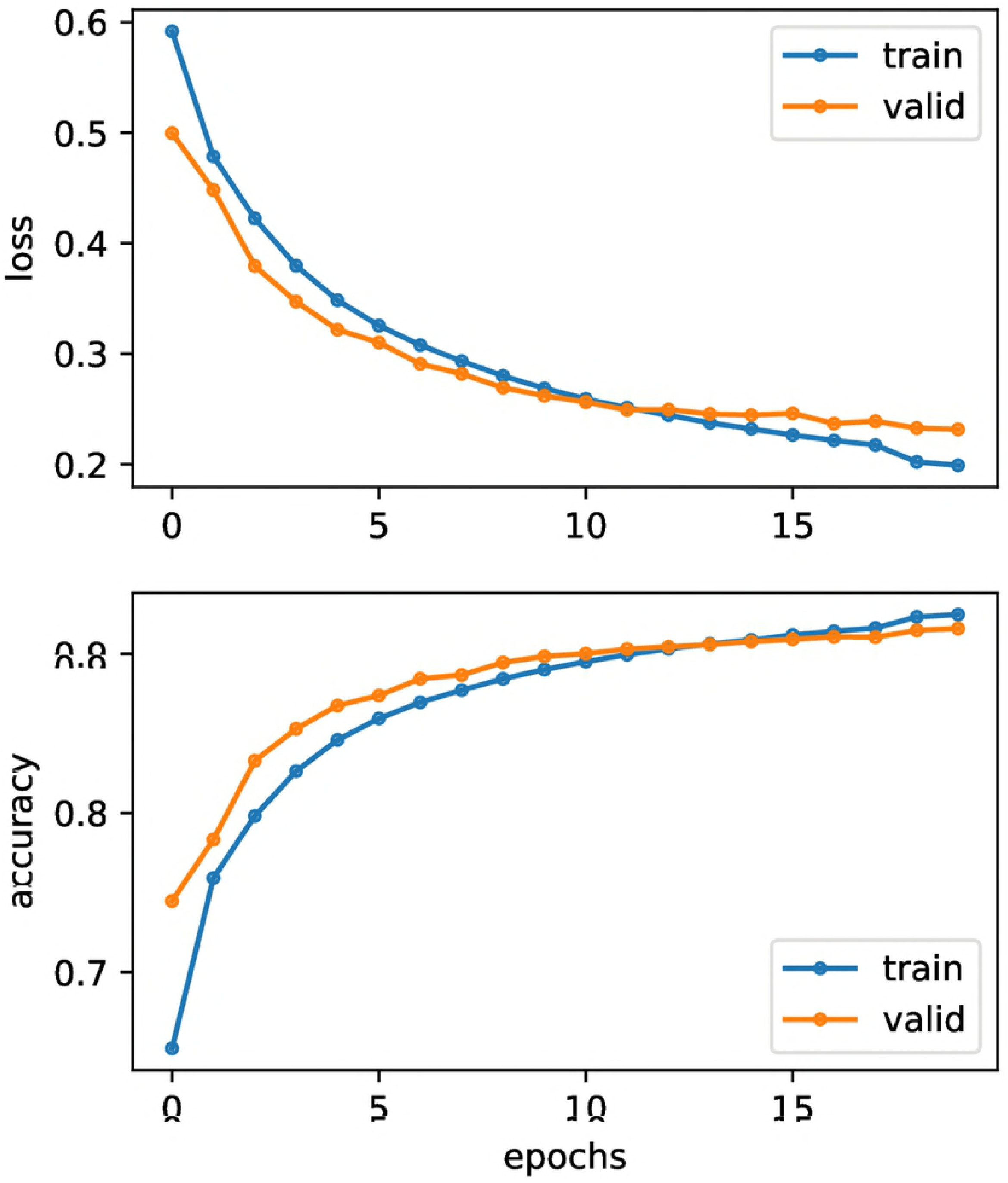
Representative example of training convergence. Loss and accuracy are shown on the top (bottom) graphs as function of epochs. Both training and validation data are balanced by oversampling the minority label.

Figs. 11 and 12 illustrate the post-training predictions for the validation data without oversampling. The distribution of scores for the positive and negative labels, Fig. 11, shows clustering for the cognate pairs at the high score value, expected for a well trained model. The achieved AUC of 0.967 (Fig. 12) suggests reliable classification.

**Fig 11.**
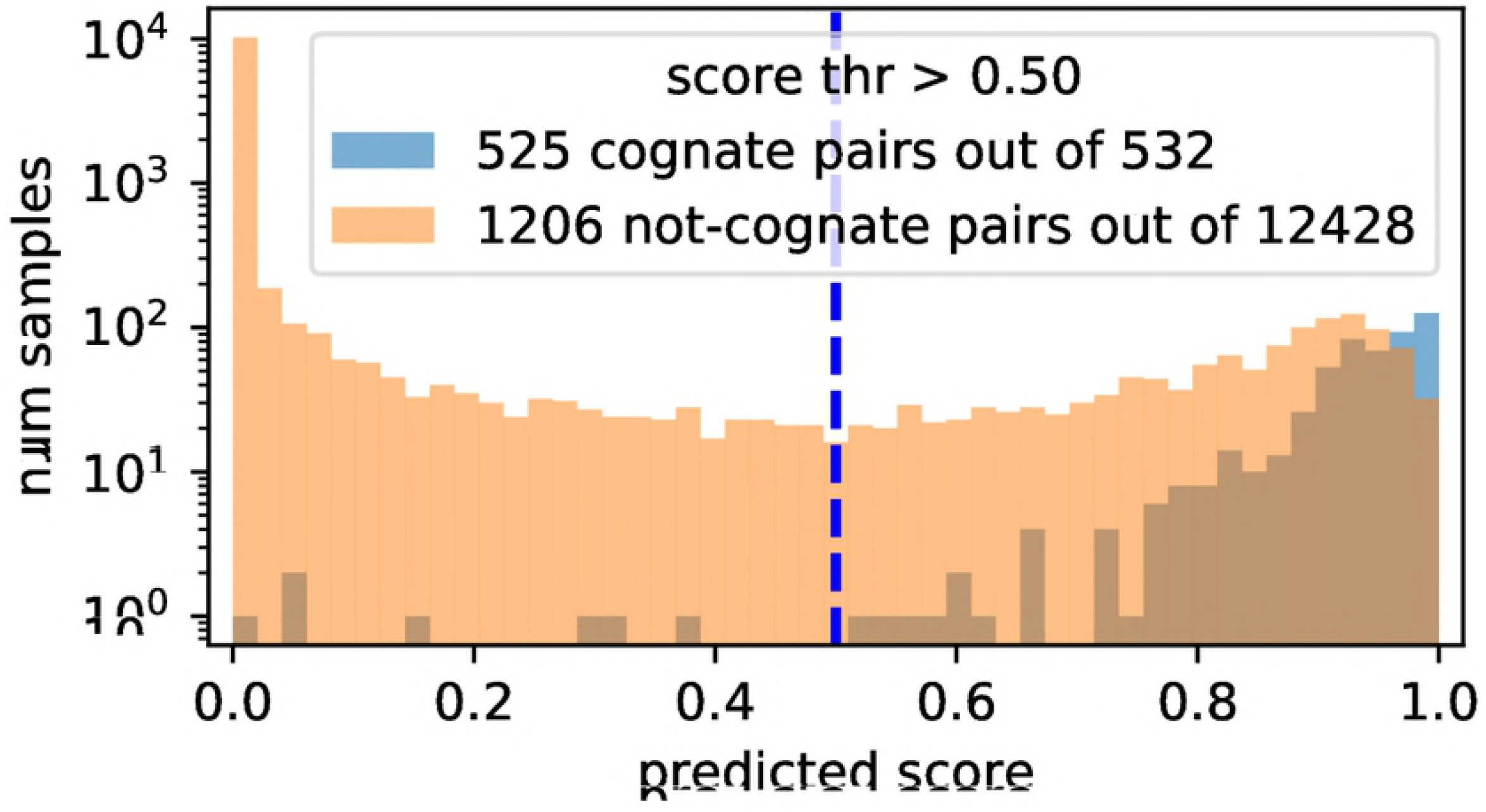
Example of post-training predictions for validation data without oversampling. The scores for positive (negative) labels are clustered around 1 (0).

**Fig 12.**
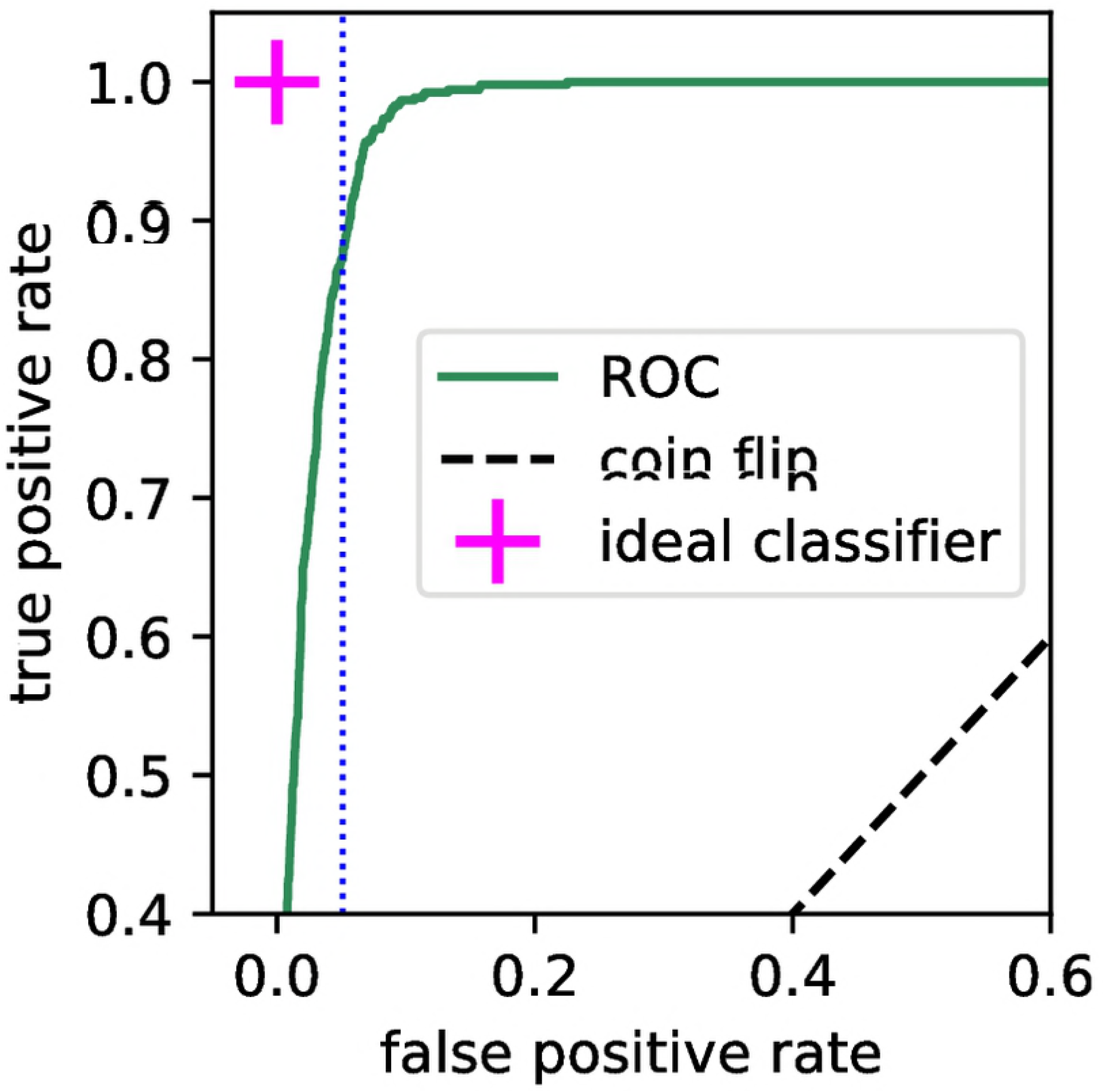
Example of ROC with AUC of 0.979 for post-training predictions of validation data without oversampling. The natural ratio of positive to negative labels (aka SNR) is 1:24. E.g. at a false positive rate of 0.05 (vertical dotted line), this model has true positive rate of 0.87, resulting in a SNR of 17. This is a 400-fold improvement over the raw input SNR.

To verify models *θ_k_* were not over-trained we computed AUC and accuracy for the k-th ‘test’ data segment not used in the *θ_k_* training. Fig. 13 shows both metrics for the 20 *θ_k_* models are stable and the spread of values is only of 0.01 that proves the models were not over-fitting the training data.

**Fig 13.**
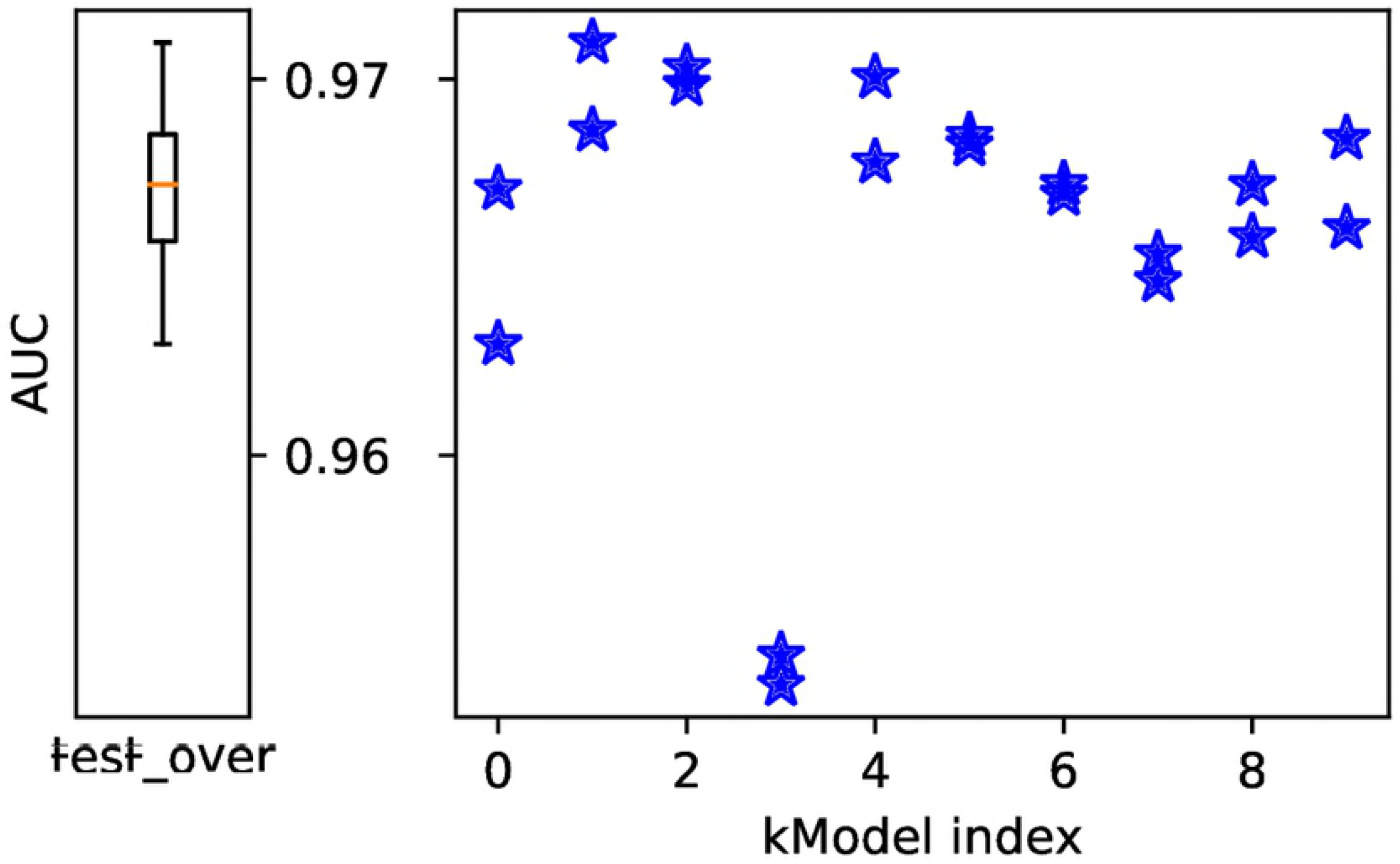
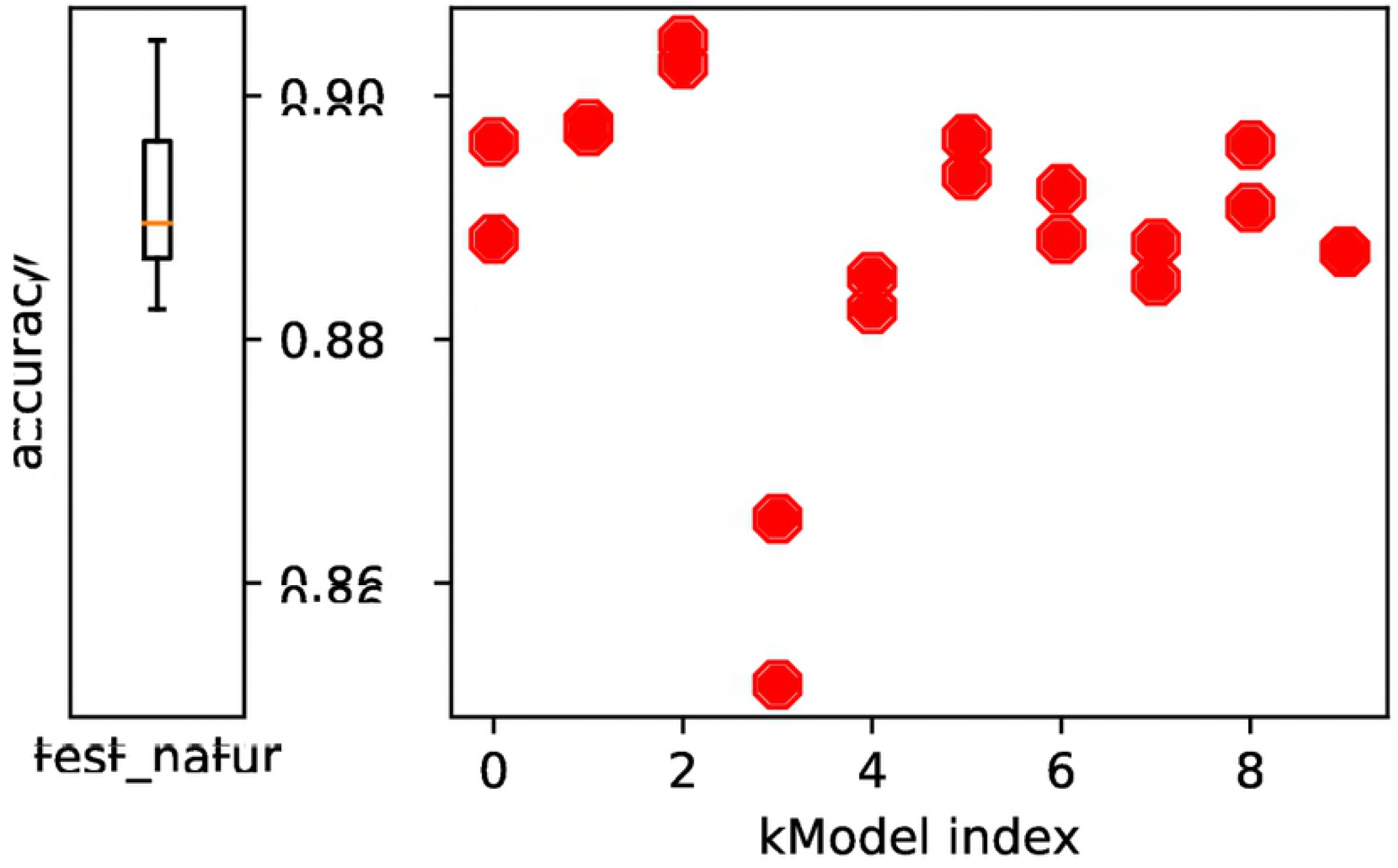
Evaluation of 20 trained models *θ_k_* on 10 ‘test’ data segments not used in the *θ_k_* training. The x-axis enumerates test segments. The box plot on the left shows the median, 25th, and 75th percentiles of the distributions. Top) AUC of ROC for imbalanced (natural) test data is 0.967. Bottom) accuracy for balanced (oversampled) test data is 0.890.

### Averaging over models

The final *ORAKLE* predictions are made by averaging scores *y_k_* from the 20 models *θ_k_*.

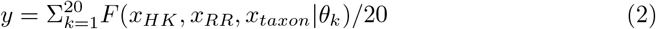

### Data and code availability

The code used in this paper and the trained model weights will be publicly available at [32] at the time of publication. The static web-page serving *ORAKLE* predictions for all 1300 species available to authors at the time of writing will be publicly available as well at at [40]. The training data can be downloaded from P2CS [19].

## Acknowledgments

This research used resources of the National Energy Research Scientific Computing Center (NERSC), a U.S. Department of Energy Office of Science User Facility operated under Contract No. DE-AC02-05CH11231.

We thank Thorsten Kurth, Mr. Prabhat, and Mustafa Mustafa from NERSC for advising in optimization of the ML code. We thank Dr. Erika Anderson for helpful edits.

## Footnotes

1 see ‘Materials and Methods’ section for details

2 Non-cognate are the mismatched pairs, see ‘Materials and Methods’ section for details.

3 If either the name of the protein or its sequence would repeat verbatim within or across species only the first occurrence was used.

## References

1. Laub MT, et al. Specificity in Two-Component Signal Transduction Pathways. Annual Review of Genetics. 2007;41(1):121–145.

2. Stock AM, et al. Two-Component Signal Transduction. Annual Review of Biochemistry. 2000;69(1):183–215.

3. Zapf J, et al. A Source of Response Regulator Autophosphatase Activity: The Critical Role of a Residue Adjacent to the Spo0F Autophosphorylation Active Site. Biochemistry. 1998;37(21):7725–7732.

4. Skerker JM, et al. Rewiring the Specificity of Two-Component Signal Transduction Systems. Cell. 2008;133(6):1043–1054.

5. Casino P, et al. Structural Insight into Partner Specificity and Phosphoryl Transfer in Two-Component Signal Transduction. Cell. 2009;139(2):325–336.

6. Li L, et al. Amino acids determining enzyme-substrate specificity in prokaryotic and eukaryotic protein kinases. Proceedings of the National Academy of Sciences. 2003;100(8):4463–4468.

7. Laub MT, et al. Phosphotransfer Profiling: Systematic Mapping of Two Component Signal Transduction Pathways and Phosphorelays. In: Two Component Signaling Systems, Part B. vol. 423 of Methods in Enzymology. Academic Press; 2007. p. 531 - 548.

8. Nan B, et al. Uncovering the Mystery of Gliding Motility in the Myxobacteria. Annual Review of Genetics. 2011;45(1):21–39.

9. Poole RK. Geobacter: The Microbe Electric’s Physiology, Ecology, and Practical Applications. In: Advances in Microbial Physiology. vol. 59 of Advances in Microbial Physiology. Academic Press; 2011. p. 1 - 100.

10. Burger L, et al. Accurate prediction of protein-protein interactions from sequence alignments using a Bayesian method. Molecular Systems Biology. 2008;4(1).

11. Black WP, et al. The orphan response regulator EpsW is a substrate of the DifE kinase and it regulates exopolysaccharide in Myxococcus xanthus. Scientific Reports. 2015;5.

12. Mike LA, et al. Two-Component System Cross-Regulation Integrates Bacillus anthracis Response to Heme and Cell Envelope Stress. PLOS Pathogens. 2014.

13. Gueudre T, et al. Simultaneous identification of specifically interacting paralogs and interprotein contacts by direct coupling analysis. Proceedings of the National Academy of Sciences. 2016;113(43):12186–12191.

14. Kara A, et al. Genome-wide prediction of prokaryotic two-component system networks using a sequence-based meta-predictor. BMC Bioinformatics. 2015;16(1):297.

15. Gers FA, et al. Learning to forget: continual prediction with LSTM. IET Conference Proceedings. 1999.

16. Nielsen H, et al. Convolutional LSTM Networks for Subcellular Localization of Proteins First Annual Danish Bioinformatics Conference Proceedings. 2015.

17. Qu Y-H, et al. On the prediction of DNA-binding proteins only from primary sequences: A deep learning approach PLoS ONE 12(12): e0188129.

18. Singh R., et al. Attend and Predict: Understanding Gene Regulation by Selective Attention on Chromatin Adv Neural Inf Process Syst. 2017 Dec; 30: 6785–6795.

19. Ortet P, et al. P2CS: updates of the prokaryotic two-component systems database, http://www.p2cs.org/. Nucleic Acids Research. 2015;43(D1):D536–D541.

20. Welch RA, et al. Extensive mosaic structure revealed by the complete genome sequence of uropathogenic Escherichia coli. Proceedings of the National Academy of Sciences. 2002;99(26):17020–17024.

21. Liu W, et al. Bacillus subtilis PhoP binds to the phoB tandem promoter exclusively within the phosphate starvation-inducible promoter. Journal of Bacteriology. 1997;179(20):6302–6310.

22. Kaiser M, et al. Nitrate repression of the Escherichia coli pfl operon is mediated by the dual sensors NarQ and NarX and the dual regulators NarL and NarP. Journal of Bacteriology. 1995;177(13):3647–55.

23. Raffa R, et al. A third envelope stress signal transduction pathway in Escherichia coli. Mol Microbiol. 2002;45(6):1599–611.

24. Bourret R, et al. Conserved aspartate residues and phosphorylation in signal transduction by the chemotaxis protein CheY. Proc Natl Acad Sci USA. 1990;87(1):51–5.

25. Townsend GE, et al. Intramolecular arrangement of sensor and regulator overcomes relaxed specificity in hybrid two-component systems. Proceedings of the National Academy of Sciences. 2013;110(2):E161–E169.

26. Loman NJ, et al. Performance comparison of benchtop high-throughput sequencing platforms. Nature Biotechnology. 2012;30:434–439.

27. Meacham F, et al. Identification and correction of systematic error in high-throughput sequence data. BMC Bioinformatics. 2011;12(1):451.

28. Abadi M, et al. A Computational Model for TensorFlow: An Introduction. In: Proceedings of the 1st ACM SIGPLAN International Workshop on Machine Learning and Programming Languages. ACM; 2017.

29. Deep Learning library for Theano and TensorFlow, https://keras.io/. 2017.

30. Good IJ. Some terminology and notation in information theory. Proceedings of the IEE - Part C: Monographs. 1956;103:200–204(4).

31. Kingma DP, et al. Adam: A Method for Stochastic Optimization. CoRR. 2014;abs/1412.6980.

32. *ORAKLE* code for predicting 2CS protein pairs with Deep Learning. https://sourceforge.net/projects/kinaseorakle

33. Ptacek J, et al. Global analysis of protein phosphorylation in yeast Nature. 2005;438:679–684

34. Deutschbauer A, et al. Evidence-Based Annotation of Gene Function in Shewanella oneidensis MR-1 Using Genome-Wide Fitness Profiling across 121 Conditions PLoS Genet 7(11): e1002385.

35. Lin WJ, et al. Threonine phosphorylation prevents promoter DNA binding of the Group B Streptococcus response regulator CovR Mol Microbiol. 2009;71:1477–95

36. Ocasio VJ, et al. Ligand-induced folding of a two-component signaling receiver domain Biochemistry. 2015;54:1353–63.

37. Burbulys D, et al. Initiation of sporulation in B. subtilis is controlled by a multicomponent phosphorelay Cell. 1991;64:545–52.

38. Wheeler RT, Shapiro L. Differential Localization of Two Histidine Kinases Controlling Bacterial Cell Differentiation Mol Cell. 1999;4:683–94.

39. Daeffler KN, et al. Engineering bacterial thiosulfate and tetrathionate sensors for detecting gut inflammation Mol Syst Biol. 2017;13:923.

40. Prediction of *ORAKLE* for 1300 species. http://portal.nersc.gov/project/nstaff/orakle2018/predictions/givenRR/

